# Assessing bite force estimates in extinct mammals and archosaurs using phylogenetic predictions

**DOI:** 10.1101/2020.11.17.386771

**Authors:** Manabu Sakamoto

## Abstract

Bite force is an ecologically important biomechanical performance measure is informative in inferring the ecology of extinct taxa. However, biomechanical modelling to estimate bite force is associated with some level of uncertainty. Here, I assess the accuracy of bite force estimates in extinct taxa using a Bayesian phylogenetic prediction model. I first fitted a phylogenetic regression model on a training set comprising extant data. The model predicts bite force from body mass and skull width while accounting for differences owning to biting position. The posterior predictive model has a 93% prediction accuracy as evaluated through leave-one-out cross-validation. I then predicted bite force in 37 species of extinct mammals and archosaurs from the posterior distribution of predictive models.

Biomechanically estimated bite forces fall within the posterior predictive distributions for all except four species of extinct taxa, and are thus as accurate as that predicted from body size and skull width, given the variation inherent in extant taxa and the amount of time available for variance to accrue. Biomechanical modelling remains a valuable means to estimate bite force in extinct taxa and should be reliably informative of functional performances and serve to provide insights into past ecologies.

## INTRODUCTION

Bite force is an ecologically important biomechanical performance measure that is also under positive phenotypic selection [1]. Therefore, biomechanical modelling of bite force is an important and informative means with which to infer the ecology of extinct taxa. However, given the lack of muscle preservation in fossil specimens, biomechanical modelling to estimate bite force is associated with some level of uncertainty. Whether this uncertainty should hinder our abilities to reliably infer biomechanical performances and past ecologies is up for debate and largely depends on the outlooks of individual researchers. Furthermore, statistical assessment of the accuracy of bite force estimates in extinct taxa has been lacking.

Here, I assess the accuracies of bite force estimates in extinct taxa using the posterior predictive distributions of a phylogenetic prediction model [2] based on bite force data in extant taxa [1], accounting for phylogenetic non-independence owing to shared ancestry [3]. Given a strong and significant relationship between bite force and predictor variables (e.g. body mass, skull widths), and phylogenetic information, it is possible to predict bite force in extinct taxa using their corresponding predictor variable values [2]. The posterior predictive distributions from such a model can serve as the null expectations. Bite forces estimated from biomechanical models independently of the predictions can then be tested against these posterior predictive distributions. If biomechanical estimates fall within the posterior predictive distribution of the model, then those estimates are as accurate as can be expected from extant data.

## MATERIAL AND METHODS

I used a Bayesian phylogenetic prediction model [2] to assess biomechanical bite force estimates in extinct taxa. Phylogenetic predictions were made from a multiple regression model of bite force (log_10_*F*_Bite_) against body mass (log_10_*M*_Body_) and skull widths (log_10_*W*_Skull_) in extant amniotes (*N*=188) [1], accounting for phylogenetic non-independence of data points owing to shared ancestry [3]. I included skull widths along with body mass as predictor variables, because the former has been shown to predict bite force accurately [4], and as the goals here are to predict bite force. I also accounted for differences in slopes amongst different groups within the data, namely bats and finches. These two clades show steeper slopes compared to the rest of the sample [1]. Additionally, I accounted for differences in bite force owing to differences in biting positions, anterior or posterior [1].

Phylogenetic predictions involve two steps. First, I fitted and evaluated a phylogenetic regression model on the training set (bite force and predictor variables in extant taxa) through Markov chain Monte Carlo (MCMC). This will produce a posterior distribution of the regression model *m*. I assessed the accuracy of this prediction model through leave-one-out cross-validation (LOOCV). LOOCV was performed by leaving one taxon out of the training set, fitting a model, and then predicting the taxon of interest using the model. I evaluated whether the predicted value differed from the observed value by calculating the proportion of the posterior predictive distribution that fell beyond the value of the biomechanical bite force estimate (*p*_MCMC_). If the biomechanical bite force estimate fell outside of the vast majority of the posterior predictive distribution (less than 5% of the posterior predictive distribution lay beyond the threshold value), then it is deemed that the biomechanical bite force value is significantly different from the posterior predictive distribution (*p*_MCMC_ < 0.05). I repeated this procedure for every tip in the phylogenetic tree over three independent MCMC chains each. Overall prediction accuracy of the phylogenetic regression model is then the number of taxa, for which the prediction is different from the observed value, out of the total number of taxa *N*=188.

I predicted bite force for the extinct taxa of interest from the posterior distribution of *m* through MCMC, given their body mass, skull widths, biting positions and phylogenetic positions. I used a phylogeny with extinct tips inserted in their relevant positions and predictions were made through MCMC so that rates of evolution along the branches leading to these extinct tips conform to Brownian motion. I then evaluated the biomechanical bite force estimates (58 estimates over 37 species; Table 1) against the posterior predictive distributions of the predictive models, using the same approach as in LOOCV. I used BayesTraits [5] for both model fitting and predicting, and R [6] for wrangling, pre-processing and post-processing of data and analytical results.

**Table 1.**
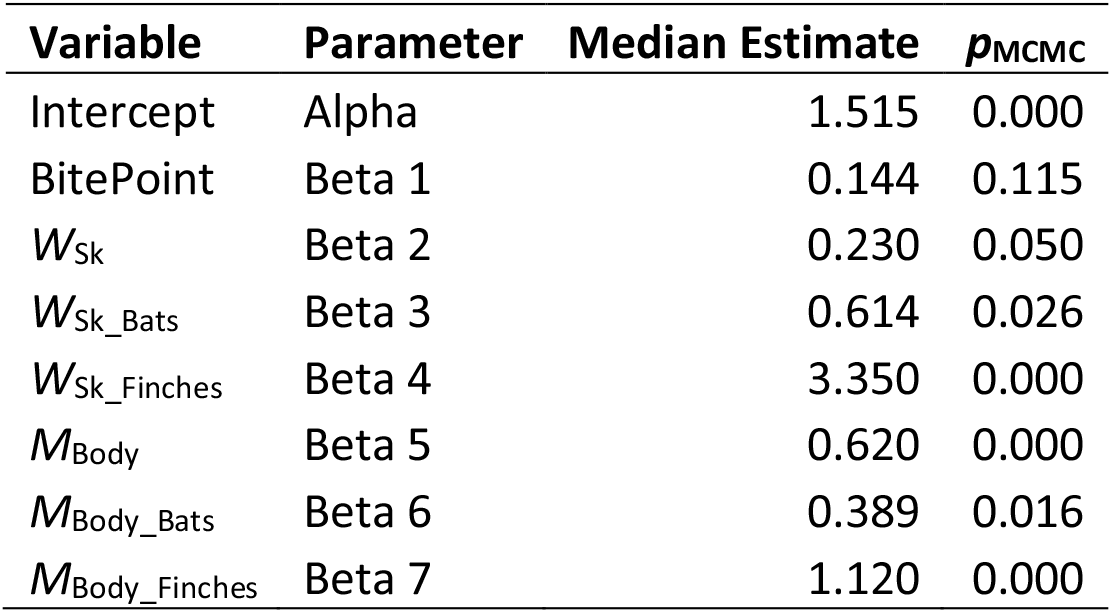
Median parameter estimates from the posterior distributions of the predictor variables and *p*_MCMC_ values.

### Comparative bite force data

I used the bite force data compiled by [1], and subset to those that also have body mass and skull widths (*N*=224; Table S1). [1] collected the bulk of the data from the literature, adding new estimates using the dry skull method [1,7].

### Phylogeny

I also used the phylogeny of [1], which is an informal supertree based on the Time Tree of Life (TTOL) [8] with fossil tips inserted manually at the appropriate phylogenetic locations [1]. Divergence times for fossil branches are based on first appearance dates (FAD) with terminal tips extended to their last appearance dates (LAD). I used the full range of temporal durations to scale the branches, as this allows for the maximum amount of time possible for trait evolution to occur [1].

## RESULTS

The phylogenetic regression model on the training set explains a high proportion of variance in bite force (*R*^2^ = 0.826). *M*_Body_ is a significant predictor in all three groups (Table 1). On the other hand, skull width (*W*_Sk_) is a significant predictor variable in bats and finches, but not in other taxa (Table 1). The effects of bite point is not significant in this model (*p*_MCMC_ = 0.115) but I include it here for subsequent predictions as this variable had significant effect in a prior study [1].

LOOCV reveals a 92.6% overall prediction accuracy for the posterior predictive model. In only 14 tips were observed values significantly different from their respective posterior predictive distributions at *p*_MCMC_ < 0.05 (Fig. 1; Table S1). These are: the jaguar, *Panthera onca*; the aardwolf, *Proteles cristatus*; 11 species of finches (including five species of Darwin’s finches, *Geospiza scandens, G. magnirostris, G. fuliginosa, Cactospiza pallida, Platyspiza crassirostris*); and the monk parakeet, *Myiopsitta monachus*.

**Figure 1.**
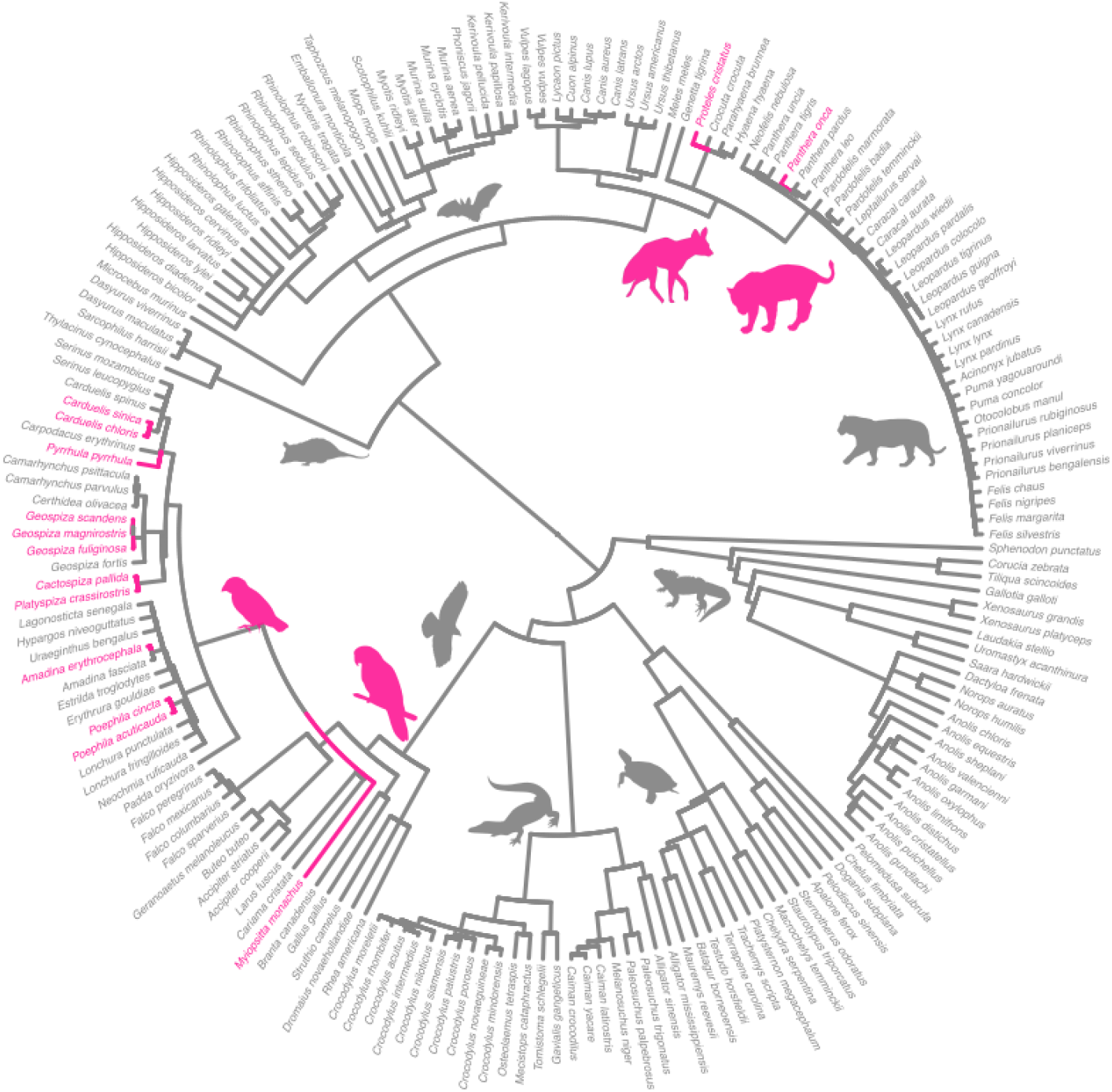
Phylogeny of extant amniotes (*N*=188) showing tips for which the observed bite force values are significantly different from the posterior predictive distributions (*p*_MCMC_ < 0.05; pink). Silhouettes from PhyloPic: *Geospiza fuliginosa*, Manabu Sakamoto, CC-BY 3.0; Psittacid, *Amazona aestiva*, Ferran-Sayol, CC0 1.0; *Panthera onca*, Manabu Sakamoto, CC-BY 3.0; *Proteles cristatus*, Margot Michaud, CC0 1.0; *Panthera tigris*, by Sarah Werning, CC-BY 3.0; Chiroptera, by Yan Wong, CC0 1.0; *Sphenodon punctatus*, by Steven Traver, CC0 1.0; *Didelphis virginiana*, Sarah Werning, CC-BY 3.0; Crocodylia, by B. Kimmel, Public Domain Mark 1.0; *Buteo buteo*, by Lauren Anderson, Public Domain Mark 1.0; *Chrysemys picta*, uncredited, Public Domain Mark 1.0.

Out of the 37 extinct taxa, four had biomechanical bite force estimates that are significantly different from the posterior predictive distributions (Table S2; Fig 2) of the phylogenetic regression model based on extant data. These are: the sabre-toothed cats, *Xenosmilus hodsonae* and *Metailurus parvulus*; the sauropodomorph dinosaur, *Plateosaurus engelhardti*; and the ornithischian dinosaur, *Stegosaurus stenops* (Fig 2). These taxa display bite forces that are significantly lower expected given their body sizes, skull widths and Brownian motion evolution.

**Figure 2.**
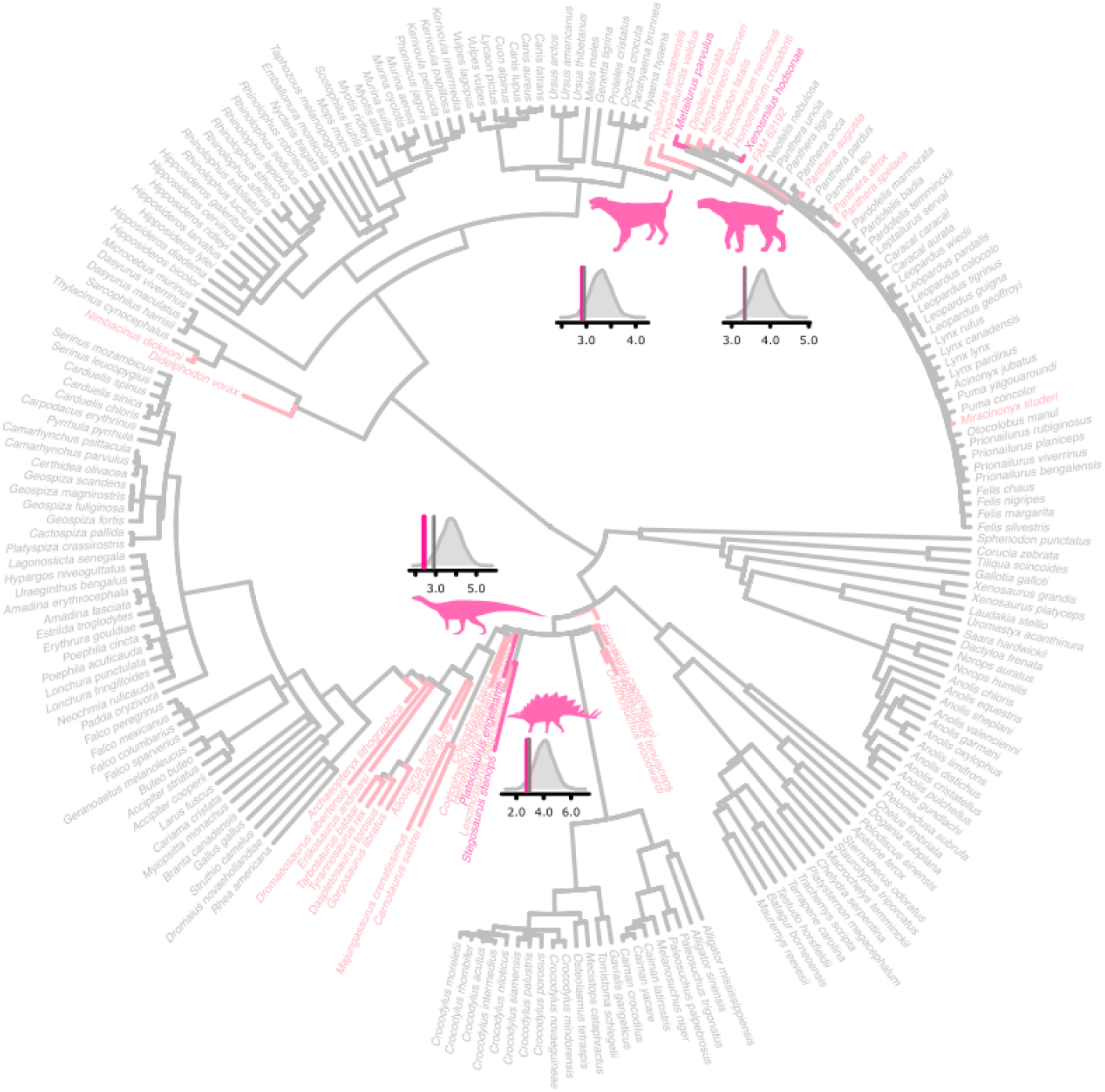
Phylogeny of amniotes showing extinct tips for which posterior predictive distributions were generated (light pink). Taxa for which observed bite forces are significantly different (*p*_MCMC_ < 0.05) from their respective posterior predictive distributions are highlighted (deep pink): *Metailurus; Xenoximulus; Plateosaurus*; and *Stegosaurus*. Posterior predictive distributions (grey) are shown with observed values (deep pink lines) and *p*_MCMC_ = 0.05 threshold values (dark grey lines) superimposed. Silhouettes from PhyloPic: *Metailurus major*, by Zimices, CC-BY 3.0; *Homotherium*, by Dantheman9758 (vectorized by T. Michael Keesey), CC-BY 3.0; *Plateosaurus*, by Andrew Knight, CC-BY 3.0; and Stegosaurus, by Andrew A. Farke, CC-BY 3.0.

## DISCUSSION

### Posterior predictive model

Overall, the posterior predictive model performs very well in predicting bite force in extant taxa (92.6% accuracy). In most taxa, bite force is as expected for their body size and skull width, under Brownian motion evolution. That is, changes in residual bite force is proportional to time and do not generally exceed expected amount of changes along the branches of the phylogenetic tree. Thus, the posterior predictive model can be used to predict bite force in extinct taxa, bracketed by extant taxa on the phylogenetic tree.

The only exceptions are in 14 taxa in which the observed bite forces are significantly different from the posterior predictive distributions (*p*_MCMC_ < 0.05; Fig 1; Table S1). These taxa were previously found to have undergone exceptional increases in rates of bite force evolution [1], indicative of positive phenotypic selection [9] on bite force. Finches in particular radiated rapidly to fill disparate ecological niches [10,11] and that their bite forces significantly deviate from those expected under Brownian motion is strongly reflective of such evolutionary processes. The jaguar is known to have more robust skulls compared to cats of similar sizes – e.g., the leopard – and have extremely strong bite forces that enable them to take on large prey. The aardwolf has extremely low bite force compared to its osteophagous relatives and this outlier status within its own family is reflected in its significant departure in bite force from expectations under Brownian motion. The monk parakeet is the only psittaciform in this dataset and its bite force clearly is not as expected given bite forces of other closely related birds (Fig 1). Thus, all significant departures from the posterior predictive distributions are consistent with our prior understanding of this dataset.

### Accuracy of bite force estimates in extinct taxa

Bite forces in extinct taxa estimated through biomechanical modelling and currently available through the literature are generally as accurate as bite forces predicted from the extant relationship between bite force and body size + skull width under Brownian motion evolution accounting for biting position. That is, biomechanically estimated bite forces in extinct taxa mostly fall within the expected range of variance for their body and skull sizes, given the variation inherent in extant data and the amount of time available for variance to accrue along the branches of the phylogenetic tree.

While the effects of accurate muscle reconstructions have been previously highlighted as a major source of discrepancies in bite force estimates (e.g., in *T. rex* between authors [12–14]), I demonstrate here that such differences are mostly negligible in a phylogenetic comparative context. At least, the variation between authors or parameterisations generally fall within expected range of variance (Figure S2). In particular, biomechanical bite force estimates for *T. rex* [1,12,13] all fall within the bulk of the posterior predictive distribution, roughly between the 50th and 75th percentiles (Fig 3).

**Figure 3.**
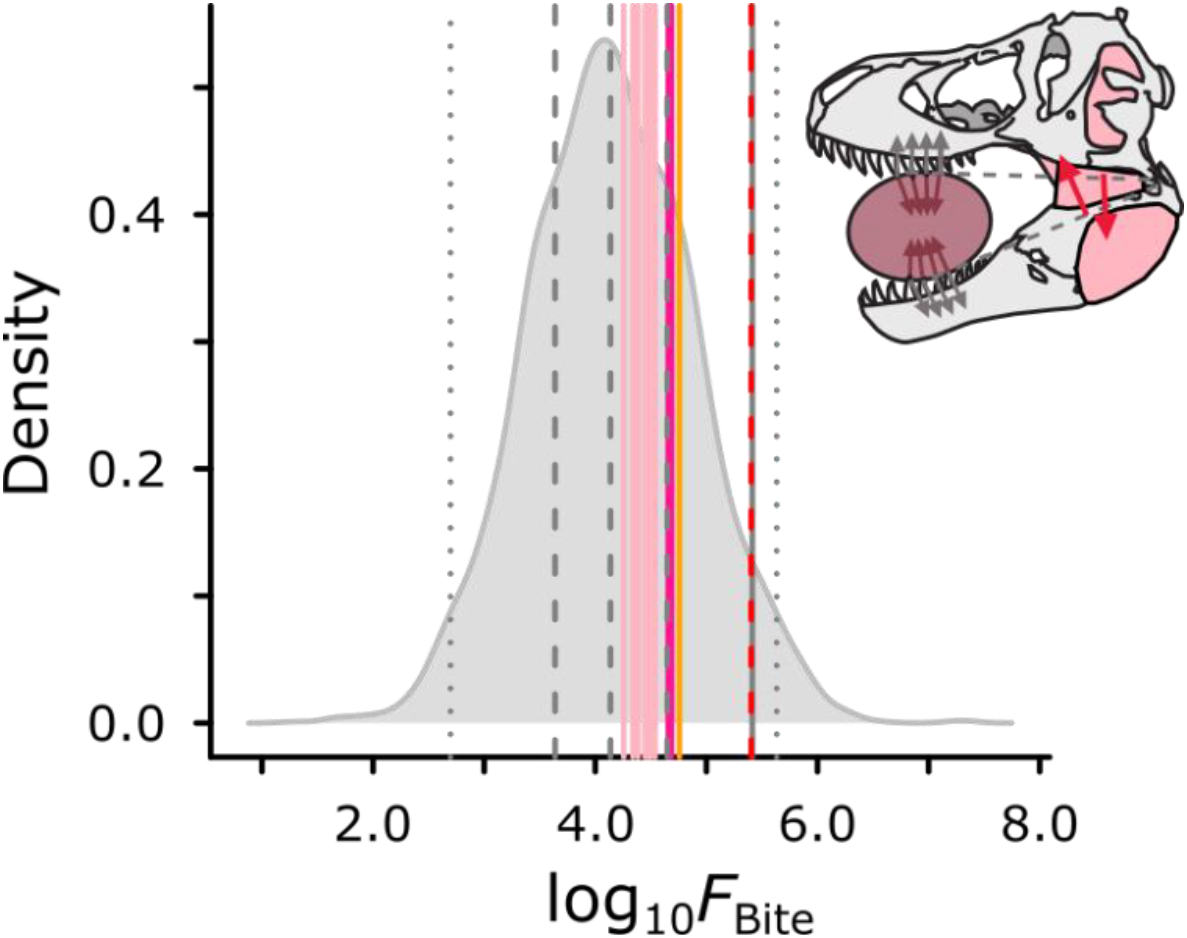
Biomechanical estimates of log_10_ bite force for *T. rex* mostly fall within the 50^th^ and 75^th^ percentiles of the posterior predictive distribution generated from a phylogenetic predictive model. Differences between studies are not statistically significant (pink, Gignac and Erickson [13]; deep pink, Sakamoto et al. [1]; orange, Falkingham and Bates [12]). Bite force extrapolated from a regression model [15] (red dashed line) sits on the threshold (solid dark grey line) at *p*_MCMC_ = 0.05. Thick dashed lines indicate 25^th^, 50^th^ and 75^th^ percentiles while tine dashed lines indicate the 2.75^th^ and 97.5^th^ percentiles.

Interestingly, a non-biomechanical bite force estimate for *T. rex* based on extrapolation of a non-phylogenetic regression model on extant data [15] can be shown here to be most likely an overestimate (*p*_MCMC_ = 0.05). Meers’s estimate [15] is higher in value than approximately 95% of the posterior predictive distribution. This is perhaps unsurprising as the extrapolated bite force of 253,123N is an order of magnitude higher than even the highest of the biomechanical estimates (57,000N [12]).

There are however exceptions to the above. Firstly, the two sabre-toothed cats, *Xenosmilus* and *Metailurus* have significantly lower bite force estimates than expected (Fig 2). Similar to the case with the extant outliers, these are entirely consistent with our prior understanding of sabre-toothed biting biomechanics [16,17]. Sabre-toothed cats are known to have smaller jaw closing muscles compared to cats of similar sizes and have been regarded as having weaker bite forces [16,18]. Indeed, bite force estimates for most sabre-toothed cats in this dataset generally fall on the lower side of the posterior predictive distributions (Fig S2). While Sakamoto et al. [1] did not find evidence for exceptional rates of bite force evolution in sabre-toothed cats using a strict threshold (>95% of rate-scaled trees and twice the background evolution [9]), they did find some evidence for elevated rates in the family Felidae, including extant conical-toothed cats, under a more relaxed threshold (>50% of rate scaled-trees). As departures from Brownian motion is here gauged through a LOOCV approach using one extinct taxon at a time, the sensitivity to detect significant departures (*p*_MCMC_ < 0.05) may be different compared to the more flexible variable-rates (VR) model [9,19] using the entire dataset of extant and extinct data. That is, once the entire range of variation is modelled, then individual departures may not stand out as exceptional rate-increases in the context of a clade exhibiting high variability in trait value. Interestingly, *Metailurus* has a superficially *Panthera*-like skull morphology, but its bite force is more reflective of sabre-toothed cats. *Metailurus* has additional biting functional morphology in line with sabre-toothed cats, such as a wider snout and larger carnassials [20].

Secondly, the two herbivorous dinosaurs, *Plateosaurus* and *Stegosaurus* have significantly lower bite forces compared to their respective posterior predictive distributions. These departures from Brownian motion are consistent with previous findings that these two taxa underwent exceptional levels of rate-increases [1]. Given that the effect of skull width is negligent in the phylogenetic regression model employed here (Table 1), the extremely small sizes of the skulls of *Plateosaurus* and *Stegosaurus* are likely not accounted for in the predictive model, and thus these taxa appear to have exceptionally low bite forces for their body sizes. As bite force estimates for herbivorous dinosaurs in general are lacking in biomechanical studies, it is difficult to say whether these extremely low values are unique to these taxa or more widespread amongst herbivorous dinosaurs.

Although the default interpretations for such outliers would be to treat them as erroneous estimates, given that outliers in extant taxa determined through LOOCV are consistently those that are known to have extreme bite forces, the same is highly likely for the extinct taxa identified here as outliers. This is especially so given the uniqueness of the outlying extinct taxa (sabre-toothed cats and herbivorous dinosaurs with extremely small heads).

### Bite force and ecological adaptations

For the most part, bite force can be explained well by body size and skull width. Bite force is known to scale strongly with body size [1] as well as skull width [4]. Skull width in particular is associated with muscle cross-sectional areas, perhaps the most influential determinant of bite force. Thus, the fact that, after accounting for these two influential variables, bite force estimates in the majority of both extant and extinct taxa fall within the expected range of residuals, offer confidence in the reliability of biomechanical methods to estimate bite force. That is, natural selection on bite force is tightly linked with body size and muscle size, and less so with residual variation. The ecological performance of bite force is predominantly associated with ecological niches dictated by size-classes. On the other hand, this means that bite force is a reliable metric for such ecologically meaningful size-classes. This is especially useful for biomechanical modelling of extinct taxa where bite force is applied as a loading parameter or simultaneously estimated.

It follows then, that outliers based on phylogenetic predictive modelling are atypical for their body size, skull width and phylogeny (Fig 1). As outliers detected here have previously been associated with elevated rates of bite force evolution [1], changes in bite force along these branches are in excess to those expected under Brownian motion evolution. Elevated rates are typically taken as evidence for positive phenotypic selection [9], but as all extinct outliers have extraordinarily low bite forces, it is more likely that selection acted on phenotypic traits that trade off with bite force. This would be gape (and clearance for hypertrophied upper canines) in sabre-toothed cats and perhaps neck elongation in *Plateosaurus* and *Stegosaurus* [21,22], which may be associated with decreases in head sizes.

## CONCLUSION

Bite force estimates in the majority of extinct taxa examined here fall within their respective posterior predictive distributions generated from a phylogenetic predictive model under Brownian motion evolution. Any discrepancies owing to uncertainties only result in deviations that are fully within the expected range of variance. On the other hand, in both extant and extinct taxa, bite force estimates are only significantly different from their respective posterior predictive distributions when such taxa are already known to have exceptionally high or low bite forces. These results combined indicate that biomechanical bite force estimates are reliable indicators/reconstructions of functional and biomechanical performances in life. This is particularly the case in the context of comparative macro-evolutionary biomechanical analyses (e.g., [1,17]), in which statistical parameters are estimated taking into account underlying evolutionary processes in the variance structure of the data.

## ACKNOWLEDGEMENTS

I would like to thank Chris Venditti for advice and guidance on phylogenetic comparative methods, Andrew Meade for support with software and high-performance cluster, and Tai Kubo for discussions.

