## Supplemental materials for "Assessing bite force estimates in extinct mammals and archosaurs using phylogenetic predictions"

Manabu Sakamoto

### Supplementary methods

#### Data

Bite force, body mass, skull width and biting positions are taken from [1]; primary sources and treatments are listed therein. The phylogenetic tree is also taken from [1]. Data were filtered to those including both body mass and skull width (Online Supplementary Materials).

#### Phylogenetic posterior predictive modelling

##### Leave-one-out cross-validation

To assess the accuracy of the prediction model, I performed leave-one-out cross-validation (LOOCV) on the extant data (*N*=188). LOOCV was performed by leaving one taxon out of the training set, fitting a model, and then predicting the taxon of interest using the model. First, I fitted a model on the training set (*N*-1) where the bite force value for the taxon of interest is replaced with a “-“ (Online Supplementary Material), which BayesTraits recognises as missing data. I used generalised least squares (GLS) through MCMC in BayesTraits to fit the phylogenetic regression model. While a variable-rates (VR) model of phenotypic evolution [2] will account for variance in bite force better, as rate heterogeneity has been observed [1], I used Brownian motion evolution for the purposes here since the goal is to generate a model that can predict any given tip from the predictors and a time-calibrated phylogeny. Model parameters are estimated on the data and tree without that tip. The posterior distribution of models is saved to disc. Second, I used this posterior distribution of models and relevant predictor variables to generate a posterior predictive distribution of bite forces for the tip of interest. For this procedure, I replaced the tip data for the taxon of interest with “?” (Online Supplementary Material), which BayesTraits recognises as the tip to estimate. For both steps I ran the chain for 350,000,000 iterations, discarding the first 250,000,000 iterations as burn-in, and then sampling every 100,000 iterations, such that the posterior distribution of models and parameters has a size of 1,000. Additional model set up and necessary BayesTraits input files can be found in the online supplementary materials. I repeated this two-step procedure for each tip on the tree.

Actual bite force values for each tip were then compared to the relevant posterior predictive distributions. If the observed value lay beyond 95% of the predicted values, then the observed value is deemed to be ‘significantly’ different from the posterior predictive distribution at *p*_MCMC_ < 0.05, where *p*_MCMC_ is the simple proportion of the posterior predictive distribution that lie on one side of the observed value. Outcomes of the LOOCV are summarised in Table S1.

##### Fitting the posterior predictive model on the training set

As LOOCV demonstrated that the posterior predictive model has high prediction accuracy (~93%), I proceeded to generate the posterior distribution of predictive models in order to predict bite force in extinct tips. This procedure requires fitting a phylogenetic regression model on the training set comprising the extant data, but bite force values for all extinct taxa replaced with “-“ (Online Supplementary Material) as in LOOCV described above. I fitted the posterior predictive model using GLS through MCMC in BayesTraits. I ran the chain for 350,000,000 iterations, discarding the first 250,000,000 iterations as burn-in, and then sampling every 100,000 iterations, such that the posterior distribution of models and parameters has a size of 1,000. The posterior sample of models was saved to file.

##### Generating posterior predictive distributions for extinct tips

I used the posterior sample of predictive models from the previous model fitting step to generate posterior predictive distributions for extinct tips based on the values of the predictor variables. I replaced bite force for the extinct taxa with “?” in the input file (Online Supplementary Material), which BayesTraits recognises as the tips to estimate. Again, I used GLS in BayesTraits, loading the sample of models along with the input and tree files. I ran the chain for 350,000,000 iterations, discarding the first 250,000,000 iterations as burn-in, and then sampling every 100,000 iterations, such that the posterior distribution of models and parameters has a size of 1,000. The output file contains the sampled estimates of the bite forces for each tip marked with “?” in the input data file (Online Supplementary Material).

##### Comparing biomechanical bite force estimates to posterior predictive distributions

I then compared published biomechanical estimates of bite force to the posterior predictive distribution generated in the previous step. Across the 37 species, I have collected 58 bite force estimates (Table S2). Some taxa are represented by multiple estimates – these could reflect independent attempts by various authors, or a range of estimates given by the same authors. As with LOOCV, I tested whether these biomechanical bite force estimates deviate from the posterior predictive distribution of bite force generated under Brownian motion evolution given the phylogenetic tree and predictor variables, represented as *p*_MCMC_ (Table S2).

**Table S1. Median predicted bite force, biomechanical bite force estimates and *p*_MCMC_ of posterior predictive distribution from LOOCV. Coloured cells are where observed values are significantly different from the corresponding posterior predictive distribution at *p*_MCMC_ < 0.05.**

| **Taxon** | **log_10_*F*_Bite_** | | ***p*_MCMC_** |
| --- | --- | --- | --- |
|  | **Pred** | **Obs** |  |
| *Felis silvestris* | 2.541 | 2.485 | 0.350 |
| *Felis margarita* | 2.354 | 2.492 | 0.159 |
| *Felis nigripes* | 2.298 | 2.313 | 0.461 |
| *Felis chaus* | 2.707 | 2.582 | 0.207 |
| *Prionailurus bengalensis* | 2.494 | 2.402 | 0.245 |
| *Prionailurus viverrinus* | 2.756 | 2.781 | 0.421 |
| *Prionailurus planiceps* | 2.389 | 2.463 | 0.295 |
| *Prionailurus rubiginosus* | 2.213 | 2.337 | 0.202 |
| *Otocolobus manul* | 2.481 | 2.492 | 0.473 |
| *Puma concolor* | 3.209 | 3.213 | 0.488 |
| *Puma yagouaroundi* | 2.616 | 2.447 | 0.144 |
| *Acinonyx jubatus* | 3.209 | 3.008 | 0.125 |
| *Lynx pardinus* | 2.735 | 2.798 | 0.341 |
| *Lynx lynx* | 2.955 | 2.823 | 0.177 |
| *Lynx canadensis* | 2.768 | 2.658 | 0.257 |
| *Lynx rufus* | 2.726 | 2.840 | 0.241 |
| *Leopardus geoffroyi* | 2.536 | 2.530 | 0.452 |
| *Leopardus guigna* | 2.298 | 2.360 | 0.117 |
| *Leopardus tigrinus* | 2.403 | 2.289 | 0.050 |
| *Leopardus colocolo* | 2.480 | 2.595 | 0.249 |
| *Leopardus pardalis* | 2.815 | 2.999 | 0.105 |
| *Leopardus wiedii* | 2.559 | 2.409 | 0.145 |
| *Caracal aurata* | 2.880 | 2.828 | 0.371 |
| *Caracal caracal* | 2.888 | 2.995 | 0.262 |
| *Leptailurus serval* | 2.854 | 2.950 | 0.308 |
| *Pardofelis temminckii* | 2.817 | 2.791 | 0.434 |
| *Pardofelis badia* | 2.507 | 2.418 | 0.284 |
| *Pardofelis marmorata* | 2.528 | 2.569 | 0.424 |
| *Panthera leo* | 3.832 | 3.607 | 0.090 |
| *Panthera pardus* | 3.337 | 3.347 | 0.474 |
| *Panthera onca* | 3.519 | 3.840 | 0.037 |
| *Panthera tigris* | 3.745 | 3.951 | 0.143 |
| *Panthera uncia* | 3.344 | 3.305 | 0.420 |
| *Neofelis nebulosa* | 3.121 | 3.199 | 0.359 |
| *Hyaena hyaena* | 3.139 | 3.250 | 0.282 |
| *Parahyaena brunnea* | 3.161 | 3.314 | 0.203 |
| *Crocuta crocuta* | 3.225 | 3.341 | 0.311 |
| *Proteles cristatus* | 2.810 | 2.352 | 0.012 |
| *Genetta tigrina* | 2.583 | 2.514 | 0.433 |
| *Meles meles* | 2.803 | 2.752 | 0.448 |
| *Ursus thibetanus* | 3.292 | 3.356 | 0.343 |
| *Ursus americanus* | 3.450 | 3.303 | 0.183 |
| *Ursus arctos* | 3.504 | 3.579 | 0.341 |
| *Canis latrans* | 2.885 | 2.835 | 0.326 |
| *Canis aureus* | 2.724 | 2.639 | 0.229 |
| *Canis lupus* | 3.001 | 3.190 | 0.075 |
| *Cuon alpinus* | 2.923 | 2.901 | 0.449 |
| *Lycaon pictus* | 3.011 | 3.047 | 0.420 |
| *Vulpes vulpes* | 2.655 | 2.679 | 0.441 |
| *Vulpes lagopus* | 2.629 | 2.599 | 0.430 |
| *Kerivoula intermedia* | 0.216 | -0.041 | 0.145 |
| *Kerivoula papillosa* | 0.655 | 0.868 | 0.172 |
| *Kerivoula pellucida* | 0.381 | 0.265 | 0.326 |
| *Phoniscus jagorii* | 0.769 | 0.808 | 0.445 |
| *Murina aenea* | 0.941 | 1.098 | 0.201 |
| *Murina cyclotis* | 1.050 | 1.076 | 0.451 |
| *Murina suilla* | 0.517 | 0.644 | 0.299 |
| *Myotis ater* | 0.350 | 0.387 | 0.441 |
| *Myotis ridleyi* | 0.266 | 0.140 | 0.282 |
| *Scotophilus kuhlii* | 1.202 | 0.963 | 0.253 |
| *Mops mops* | 1.385 | 0.849 | 0.130 |
| *Taphozous melanopogon* | 0.901 | 0.891 | 0.489 |
| *Emballonura monticola* | 0.247 | 0.025 | 0.319 |
| *Nycteris tragata* | 0.866 | 0.975 | 0.412 |
| *Rhinolophus robinsoni* | 0.502 | 0.344 | 0.296 |
| *Rhinolophus sedulus* | 0.342 | 0.504 | 0.295 |
| *Rhinolophus lepidus* | 0.299 | 0.248 | 0.427 |
| *Rhinolophus stheno* | 0.387 | 0.455 | 0.391 |
| *Rhinolophus affinis* | 0.739 | 0.638 | 0.346 |
| *Rhinolophus trifoliatus* | 0.723 | 0.823 | 0.387 |
| *Rhinolophus luctus* | 1.072 | 1.260 | 0.302 |
| *Hipposideros galeritus* | 0.342 | 0.053 | 0.212 |
| *Hipposideros cervinus* | 0.149 | 0.207 | 0.435 |
| *Hipposideros ridleyi* | 0.499 | 0.573 | 0.423 |
| *Hipposideros larvatus* | 0.800 | 0.973 | 0.294 |
| *Hipposideros lylei* | 1.331 | 1.212 | 0.363 |
| *Hipposideros diadema* | 1.430 | 1.395 | 0.456 |
| *Hipposideros bicolor* | 0.449 | 0.560 | 0.387 |
| *Microcebus murinus* | 1.423 | 1.552 | 0.428 |
| *Dasyurus viverrinus* | 2.197 | 2.181 | 0.461 |
| *Dasyurus maculatus* | 2.525 | 2.580 | 0.373 |
| *Sarcophilus harrisii* | 2.897 | 2.849 | 0.390 |
| *Thylacinus cynocephalus* | 3.314 | 3.161 | 0.379 |
| *Serinus mozambicus* | 0.284 | 0.477 | 0.162 |
| *Serinus leucopygius* | 0.105 | 0.301 | 0.176 |
| *Carduelis spinus* | 0.431 | 0.477 | 0.401 |
| *Carduelis sinica* | 0.664 | 0.903 | 0.027 |
| *Carduelis chloris* | 1.450 | 1.146 | 0.005 |
| *Carpodacus erythrinus* | 0.910 | 0.778 | 0.307 |
| *Pyrrhula pyrrhula* | 1.125 | 0.699 | 0.046 |
| *Camarhynchus psittacula* | 0.979 | 1.003 | 0.298 |
| *Camarhynchus parvulus* | 0.668 | 0.662 | 0.444 |
| *Certhidea olivacea* | 0.306 | 0.223 | 0.245 |
| *Geospiza scandens* | 1.198 | 0.890 | 0.000 |
| *Geospiza magnirostris* | 1.437 | 1.850 | 0.000 |
| *Geospiza fuliginosa* | 0.468 | 0.721 | 0.000 |
| *Geospiza fortis* | 1.133 | 1.241 | 0.305 |
| *Cactospiza pallida* | 0.828 | 0.201 | 0.000 |
| *Platyspiza crassirostris* | 0.630 | 1.192 | 0.000 |
| *Lagonosticta senegala* | -0.313 | 0.000 | 0.060 |
| *Hypargos niveoguttatus* | 0.684 | 0.477 | 0.155 |
| *Uraeginthus bengalus* | 0.150 | 0.000 | 0.227 |
| *Amadina erythrocephala* | 0.840 | 0.602 | 0.016 |
| *Amadina fasciata* | 0.583 | 0.699 | 0.148 |
| *Estrilda troglodytes* | -0.301 | 0.000 | 0.085 |
| *Erythrura gouldiae* | 0.522 | 0.602 | 0.358 |
| *Poephila cincta* | 0.942 | 0.477 | 0.000 |
| *Poephila acuticauda* | -0.032 | 0.477 | 0.000 |
| *Lonchura punctulata* | 0.408 | 0.602 | 0.194 |
| *Lonchura fringilloides* | 0.725 | 0.699 | 0.452 |
| *Neochmia ruficauda* | 0.351 | 0.301 | 0.404 |
| *Padda oryzivora* | 1.307 | 1.000 | 0.209 |
| *Falco peregrinus* | 1.141 | 1.113 | 0.431 |
| *Falco mexicanus* | 1.030 | 1.140 | 0.254 |
| *Falco columbarius* | 0.732 | 0.676 | 0.380 |
| *Falco sparverius* | 0.573 | 0.601 | 0.457 |
| *Geranoaetus melanoleucus* | 1.319 | 1.699 | 0.060 |
| *Buteo buteo* | 1.092 | 0.947 | 0.273 |
| *Accipiter striatus* | 0.396 | 0.152 | 0.213 |
| *Accipiter cooperii* | 0.650 | 0.491 | 0.303 |
| *Larus fuscus* | 0.868 | 1.171 | 0.296 |
| *Cariama cristata* | 1.046 | 1.288 | 0.340 |
| *Myiopsitta monachus* | 0.216 | 1.224 | 0.040 |
| *Branta canadensis* | 1.266 | 1.440 | 0.391 |
| *Gallus gallus* | 0.927 | 0.944 | 0.486 |
| *Struthio camelus* | 1.904 | 1.722 | 0.374 |
| *Dromaius novaehollandiae* | 1.699 | 0.964 | 0.096 |
| *Rhea americana* | 1.394 | 1.412 | 0.486 |
| *Crocodylus moreletii* | 3.446 | 3.643 | 0.081 |
| *Crocodylus rhombifer* | 3.326 | 3.324 | 0.492 |
| *Crocodylus acutus* | 3.760 | 3.602 | 0.068 |
| *Crocodylus intermedius* | 3.713 | 3.798 | 0.198 |
| *Crocodylus niloticus* | 3.693 | 3.483 | 0.135 |
| *Crocodylus siamensis* | 3.399 | 3.533 | 0.236 |
| *Crocodylus palustris* | 3.905 | 3.863 | 0.405 |
| *Crocodylus porosus* | 3.978 | 3.953 | 0.458 |
| *Crocodylus novaeguineae* | 3.748 | 3.729 | 0.451 |
| *Crocodylus mindorensis* | 3.399 | 3.437 | 0.410 |
| *Osteolaemus tetraspis* | 2.988 | 3.252 | 0.185 |
| *Mecistops cataphractus* | 3.528 | 3.318 | 0.239 |
| *Tomistoma schlegelii* | 3.451 | 3.531 | 0.404 |
| *Gavialis gangeticus* | 3.521 | 3.278 | 0.236 |
| *Caiman crocodilus* | 3.003 | 3.085 | 0.278 |
| *Caiman yacare* | 3.081 | 2.987 | 0.259 |
| *Caiman latirostris* | 3.155 | 3.166 | 0.475 |
| *Melanosuchus niger* | 3.427 | 3.431 | 0.505 |
| *Paleosuchus palpebrosus* | 2.962 | 2.954 | 0.488 |
| *Paleosuchus trigonatus* | 3.043 | 3.034 | 0.487 |
| *Alligator sinensis* | 2.922 | 3.035 | 0.413 |
| *Alligator mississippiensis* | 3.723 | 3.709 | 0.486 |
| *Mauremys reevesii* | 1.095 | 1.301 | 0.330 |
| *Batagur borneoensis* | 2.533 | 2.167 | 0.219 |
| *Testudo horsfieldii* | 1.550 | 1.255 | 0.287 |
| *Terrapene carolina* | 1.424 | 1.398 | 0.479 |
| *Trachemys scripta* | 1.334 | 1.176 | 0.366 |
| *Platysternon megacephalum* | 1.211 | 1.623 | 0.222 |
| *Chelydra serpentina* | 2.607 | 2.320 | 0.263 |
| *Macrochelys temminckii* | 1.649 | 2.199 | 0.122 |
| *Staurotypus triporcatus* | 1.729 | 2.143 | 0.216 |
| *Sternotherus odoratus* | 1.746 | 1.491 | 0.324 |
| *Apalone ferox* | 1.315 | 1.623 | 0.282 |
| *Pelodiscus sinensis* | 1.722 | 1.771 | 0.466 |
| *Dogania subplana* | 1.757 | 1.568 | 0.379 |
| *Chelus fimbriata* | 1.425 | 0.699 | 0.192 |
| *Pelomedusa subrufa* | 0.968 | 0.903 | 0.472 |
| *Anolis gundlachi* | 0.813 | 0.823 | 0.485 |
| *Anolis pulchellus* | 0.383 | 0.401 | 0.473 |
| *Anolis cristatellus* | 0.861 | 0.877 | 0.482 |
| *Anolis distichus* | 0.600 | 0.447 | 0.325 |
| *Anolis limifrons* | 0.356 | 0.292 | 0.396 |
| *Anolis oxylophus* | 0.634 | 0.646 | 0.477 |
| *Anolis garmani* | 1.073 | 1.043 | 0.452 |
| *Anolis valencienni* | 0.755 | 0.826 | 0.409 |
| *Anolis sheplani* | 0.114 | 0.009 | 0.394 |
| *Anolis equestris* | 1.488 | 1.761 | 0.239 |
| *Anolis chloris* | 0.441 | 0.768 | 0.231 |
| *Norops humilis* | 0.450 | 0.328 | 0.368 |
| *Norops auratus* | 0.424 | 0.458 | 0.461 |
| *Dactyloa frenata* | 1.452 | 1.436 | 0.487 |
| *Saara hardwickii* | 1.677 | 1.708 | 0.481 |
| *Uromastyx acanthinura* | 1.782 | 1.772 | 0.489 |
| *Laudakia stellio* | 1.222 | 1.009 | 0.384 |
| *Xenosaurus platyceps* | 1.357 | 1.094 | 0.214 |
| *Xenosaurus grandis* | 1.011 | 1.277 | 0.213 |
| *Gallotia galloti* | 1.204 | 2.036 | 0.163 |
| *Tiliqua scincoides* | 1.880 | 1.616 | 0.292 |
| *Corucia zebrata* | 1.616 | 1.770 | 0.366 |
| *Sphenodon punctatus* | 1.910 | 2.377 | 0.315 |

**Table S2. Median predicted bite force, biomechanical bite force estimates and *p*_MCMC_ of posterior predictive distribution in extinct taxa**

| **Taxon** | **log_10_*F*_Bite_** | | ***p*_MCMC_** | **Source** | **Additional info** |
| --- | --- | --- | --- | --- | --- |
|  | **Pred** | **Obs** |  |  |  |
| *Miracinonyx studeri* | 3.049 | 2.983 | 0.295 | [1] |  |
| *Panthera spelaea* | 3.675 | 3.572 | 0.191 | [1] |  |
| *Panthera atrox* | 3.889 | 3.702 | 0.053 | [1] |  |
| *Panthera augusta* | 3.797 | 3.630 | 0.086 | [1] |  |
| *FAM 62192* | 3.181 | 3.113 | 0.338 | [1] | FAM 62192 |
| *Xenosmilus hodsonae* | 3.792 | 3.326 | 0.037 | [1] |  |
| *Homotherium crusafonti* | 3.557 | 3.224 | 0.078 | [1] |  |
| *Homotherium nestianus* | 3.792 | 3.411 | 0.065 | [1] |  |
| *Smilodon fatalis* | 3.729 | 3.377 | 0.074 | [3] |  |
| *Smilodon fatalis* | 3.729 | 3.500 | 0.183 | [1] |  |
| *Megantereon falconeri* | 3.406 | 3.077 | 0.085 | [1] |  |
| *Dinofelis cristata* | 3.577 | 3.302 | 0.125 | [1] |  |
| *Metailurus parvulus* | 3.296 | 2.907 | 0.032 | [1] |  |
| *Hyperailurictis validus* | 3.225 | 3.267 | 0.432 | [1] |  |
| *Proailurus lemanensis* | 2.973 | 2.985 | 0.484 | [1] | MNHN-1903-20 |
| *Nimbacinus dicksoni* | 2.617 | 2.759 | 0.281 | [3] |  |
| *Didelphodon vorax* | 2.421 | 2.452 | 0.476 | [4] | NDGS 431 |
| *Archaeopteryx lithographica* | 0.962 | 0.490 | 0.139 | [1] | Mayr 2007; Paul 1988 |
| *Dromaeosaurus albertensis* | 2.212 | 2.947 | 0.150 | [1] | AMNH.5356 |
| *Erlikosaurus andrewsi* | 3.090 | 2.428 | 0.154 | [5] |  |
| *Erlikosaurus andrewsi* | 3.090 | 2.384 | 0.138 | [6] | IGM 100/111 |
| *Erlikosaurus andrewsi* | 3.090 | 2.334 | 0.122 | [6] | IGM 100/111 |
| *Erlikosaurus andrewsi* | 3.090 | 2.425 | 0.153 | [6] | IGM 100/111 |
| *Tarbosaurus bataar* | 3.760 | 4.385 | 0.204 | [1] | Witmer pers comms |
| *Tyrannosaurus rex* | 4.136 | 4.757 | 0.212 | [7] | BHI 3033 |
| *Tyrannosaurus rex* | 4.136 | 4.657 | 0.246 | [1] | BHI 3033 |
| *Tyrannosaurus rex* | 4.136 | 4.686 | 0.235 | [1] | AMNH5027 |
| *Tyrannosaurus rex* | 4.136 | 4.538 | 0.305 | [8] | FMNH PR 2081 |
| *Tyrannosaurus rex* | 4.136 | 4.495 | 0.322 | [8] | LACM 23844 |
| *Tyrannosaurus rex* | 4.136 | 4.484 | 0.326 | [8] | MOR 980 |
| *Tyrannosaurus rex* | 4.136 | 4.449 | 0.341 | [8] | MOR 008 |
| *Tyrannosaurus rex* | 4.136 | 4.385 | 0.372 | [8] | BHI 3033 |
| *Tyrannosaurus rex* | 4.136 | 4.338 | 0.390 | [8] | RTMP 81.6.1 |
| *Tyrannosaurus rex* | 4.136 | 4.256 | 0.429 | [8] | BHI 4100 |
| *Daspletosaurus torosus* | 3.881 | 4.210 | 0.325 | [1] | FMNH PR308 |
| *Daspletosaurus torosus* | 3.881 | 4.221 | 0.319 | [1] | NMC 8506 |
| *Gorgosaurus libratus* | 3.766 | 4.140 | 0.305 | [1] | USNM 12814 |
| *Allosaurus fragilis* | 3.596 | 3.941 | 0.266 | [7] | SMA 0005 |
| *Allosaurus fragilis* | 3.596 | 3.973 | 0.249 | [1] | Madsen1976 |
| *Allosaurus fragilis* | 3.596 | 3.553 | 0.470 | [9] | DNM 2560 |
| *Sinraptor dongi* | 3.622 | 4.035 | 0.223 | [1] | IVPP.10600 |
| *Majungasaurus crenatissimus* | 3.744 | 3.895 | 0.424 | [1] | FMNH.PR2100 |
| *Carnotaurus sastrei* | 3.831 | 3.722 | 0.449 | [10] |  |
| *Coelophysis bauri* | 2.515 | 2.460 | 0.458 | [1] | Colbert 1989 |
| *Coelophysis rhodesiensis* | 2.545 | 2.594 | 0.463 | [1] | Colbert 1989 |
| *Herrerasaurus ischigualastensis* | 3.025 | 3.287 | 0.254 | [1] | PVSJ407 |
| *Plateosaurus engelhardti* | 3.759 | 2.441 | 0.004 | [6] | MB.R.1937 |
| *Plateosaurus engelhardti* | 3.759 | 2.450 | 0.004 | [6] | MB.R.1937 |
| *Plateosaurus engelhardti* | 3.759 | 2.384 | 0.003 | [6] | MB.R.1937 |
| *Lesothosaurus diagnosticus* | 2.342 | 2.397 | 0.455 | [1] | Sereno 1991a/Nesbitt 2011 |
| *Stegosaurus stenops* | 4.054 | 2.740 | 0.027 | [11] |  |
| *Stegosaurus stenops* | 4.054 | 2.914 | 0.044 | [6] | NHMUK PV R36730 |
| *Stegosaurus stenops* | 4.054 | 2.854 | 0.036 | [6] | NHMUK PV R36730 |
| *Stegosaurus stenops* | 4.054 | 2.949 | 0.048 | [6] | NHMUK PV R36730 |
| *Ornithosuchus woodwardi* | 3.567 | 3.854 | 0.279 | [1] | Sereno 1991b/Walker 1964 |
| *Riojasuchus tenuisceps* | 2.570 | 2.365 | 0.343 | [1] | Sereno 1991b/Nesbitt 2011 |
| *Parasuchus hislopi* | 3.384 | 3.292 | 0.412 | [1] | Sereno 1991b/Chatterjee 1978 |
| *Euparkeria capensis* | 1.784 | 2.334 | 0.062 | [1] | Ewer 1965 |
